# U2 snRNP recognizes the branch site through a loaded-spring strand-invasion mechanism

**DOI:** 10.1101/2025.10.31.685824

**Authors:** Pavlína Pokorná, Vlad Pena, Alessandra Magistrato

## Abstract

Recognition of the branch sequence (BS) by U2 snRNP is a pivotal step in pre-mRNA splicing and spliceosome assembly. Structural studies suggest that BS recognition occurs through a toe-hold-mediated strand-invasion mechanism, in which U2 snRNA progressively base-pairs with the intron to form the branch helix. However, given the limited complementarity between U2 snRNA and the intronic BS, it remains unclear how such strand invasion can occur spontaneously. Here, using all-atom and coarse-grained molecular dynamics simulations, we show that strand invasion proceeds spontaneously once the toehold region is engaged and the TAT-SF1 factor is released. The key finding is that the branch-stem loop (BSL) of U2 snRNA is maintained in a supercoiled, high-energy conformation by TAT-SF1, which acts as a molecular latch holding the BSL in a poised “loaded-spring” state. Displacement of TAT-SF1 allows the BSL to relax, releasing the stored conformational energy that drives strand invasion through local strand-slip and base-pair exchange. Moreover, the simulations reveal that strand invasion can proceed bidirectionally, refining previous models of U2–BS pairing. This work establishes a loaded-spring mechanism as a key physical driver of toehold-mediated strand invasion underlying branch-site recognition within early spliceosome.

## INTRODUCTION

Strand invasion, also known as strand displacement, occurs when a single-stranded DNA or RNA strand invades a duplex, displacing one of its strands to hybridize with the other. This process occurs without external protein drivers, such as helicases [1]. *In vitro*, strand invasion processes find applications in nanotechnology [2], [3]. *In vivo*, this mechanism is suggested to mediate nucleic acid rearrangements and recognition, being implicated in paramount biological processes such as transcription regulation, protein synthesis and pre-mRNA splicing [1].

Here we aimed to resolve the molecular mechanism of strand displacement in the context of pre-mRNA splicing. Splicing consists of removing non-coding regions (introns) and ligating the coding regions (exons) to produce functional protein-coding RNA and long non-coding RNAs [4], [5]. Splicing is orchestrated by a spliceosome, large and dynamic ribonucleoprotein (RNP) engine [6], [7], [8], [9]. Splicing fidelity, critical to maintain proteome integrity, relies on accurate recognition of specific pre-mRNA sequences [10]. These are the exon-intron boundaries (5′- and 3′-splice site) and the branch sequence (BS). The BS contains the branch point adenosine (BPA), which later initiates the first catalytic step of the splicing reaction [11], [12]. During the spliceosome assembly, the BS is initially bound by the SF1 protein and, subsequently, handed over to U2 small nuclear (sn)RNP, where it pairs the U2 snRNA and contacts the SF3b complex [13], [14].

Based on cryo-EM studies, Cretu and coworkers have proposed that branch site recognition by U2 snRNP occurs stepwise, through an intron-mediated strand displacement [15]. Initially, upon binding to U2 snRNA/SF3b complex, the intron BS hybridizes with a branch stem-loop (BSL) of U2 snRNA (**Figure 1**) [16]. Finally, the pairing extends by forming a full branch helix, where the BPA is bulged out [13], [17], [18], [19], [20].

**Figure 1:**
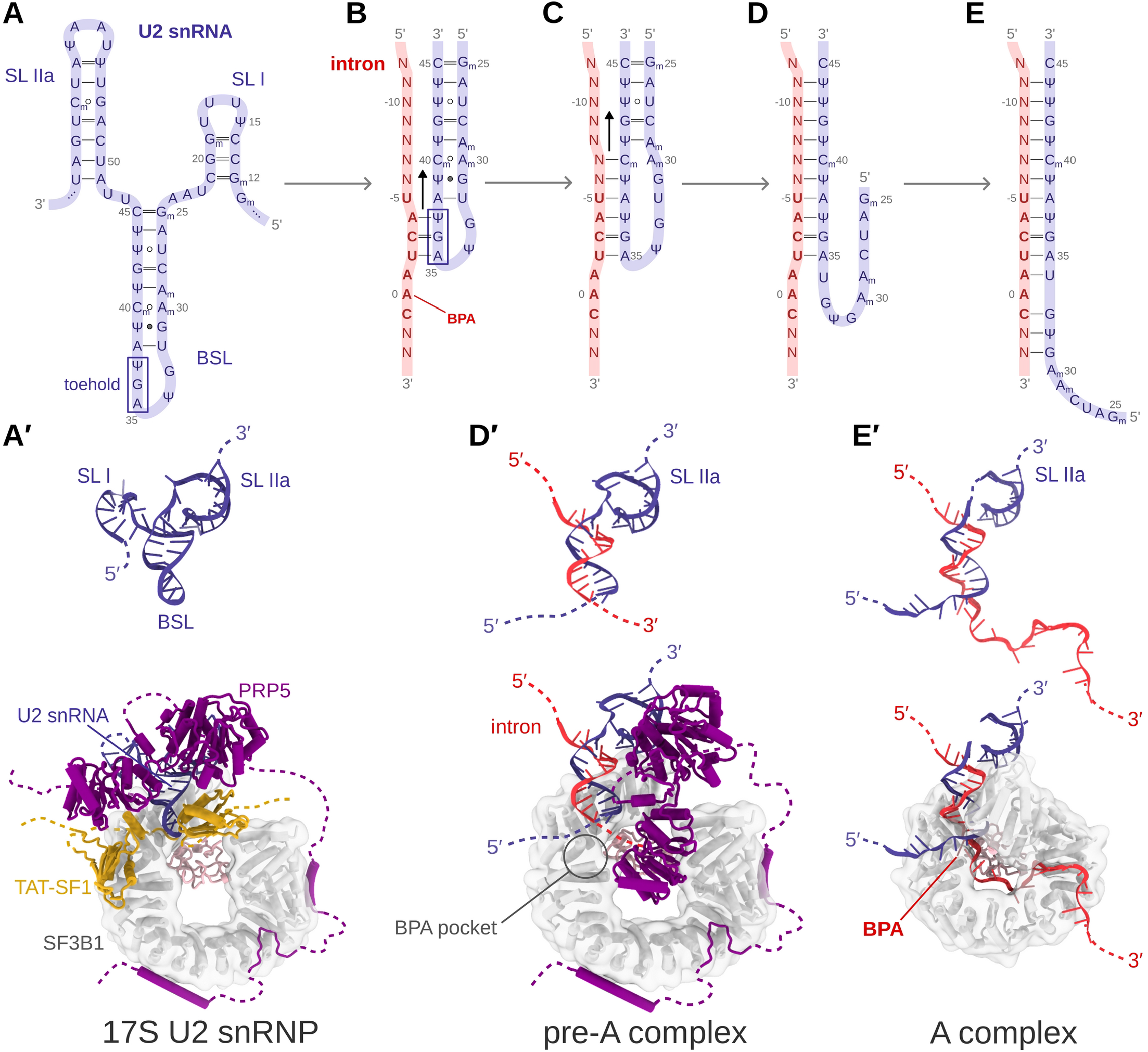
Strand displacement mechanism as proposed by Cretu et al. [15]. **(A)** Scheme of 3′ terminal portion of U2 snRNA. For clarity, stem loops SL I and SL IIa are omitted in panels (B)-(E). **(B)** Hypothesized formation of the initial toehold between the intron (red) and the BSL of U2 snRNA (blue). The yeast consensus BS sequence 5′-UACUA<u>A</u>C-3′ is marked with bold red letters. The corresponding human consensus motif is 5′-YNYUR<u>A</u>Y-3′, where <u>A</u> is the BPA, Y stands for pyrimidine, R for purine, and N for any base [4], [23]. **(C)** Hypothesized growth mechanism of the branch helix. The 5’-part of the intron establishes additional base pair interactions with the BSL 3′-end. Notably, the intermediates depicted in **(B)** and **(C)** were not captured experimentally. **(D)** The invasion of the intron strand continues with the formation of the branch helix, while the 5′-end of the BSL dissociates. **(E)** Complete formation of the branch helix. **(A′), (D′)**, and **(E′)** show RNA 3D structures corresponding those sketched in **(A), (D)**, and **(E)**. These 3D structures are cut from cryo-EM structures of the states captured in PDBs 7EVO [14], 7VPX [14], and 6FF4 [17]), respectively. The corresponding models, embedded in the SF3b complex, are shown in the bottom. Here the SF3B1, PRP5 and TAT-SF1 proteins are shown as white, magenta and yellow cartoons.

In more detail, cryo-EM structures of 17S U2 snRNP showed that three bases are extruded from the loop when the BSL binds to the SF3B1 protein - the main component of the SF3b complex. The rest of the BSL is instead masked by the TAT-SF1 protein (**Figure 1A**). Subsequent TAT-SF1 removal and intron binding to BSL is promoted by PRP5 (also known as DDX46) ATPase/helicase, which also exerts intron proofreading [13], [16], [21], [22]. Cryo-EM structure of a spliceosome stalled with spliceostatin A has revealed an intermediate conformation, found half-way through the branch duplex formation. This is a pre-A spliceosome, captured after PRP5 action, where U2 snRNA has paired the upstream (5’ direction) sequence of BS to form a precursor of the branch helix (**Figure 1D**) [14], [15]. Finally, BS recognition ends with complete formation of the branch helix, which is embraced by SF3B1 HEAT-like solenoid structure and the BPA docks into a binding pocket at the interface of the SF3B1/PHF5A proteins (**Figure 1E**). The proposed branch helix formation mechanism [15] starts with the three extruded bases of the BSL initially base pairing with the intron. Thus, these bases serve as a toehold that anchors the BSL to the intron. The subsequent growth of the branch helix would occur via strand displacement in which the intron is expected to replace the 5′-end of BSL (**Figure 1**). This results in a complete opening of BSL [15], [16]. However, strand displacement typically involves good base pair complementarity to energetically drive the strand migration [1]. Therefore, given that introns typically exhibit limited base pair complementarity with U2 snRNA, it remains unclear how U2 snRNA can recognize the branch sequence via a strand invasion mechanism, especially since the branch sequence possesses solely a short, conserved motif that shows a significant degree of degeneracy [4], [23] (**Figure 1**).

Here, we combined all-atom and coarse-grained molecular dynamics (MD) simulations to elucidate how U2 snRNA recognizes the branch sequence through strand invasion despite weak complementarity. We show that release of the TAT-SF1 protein triggers relaxation of a supercoiled, high-energy conformation of the U2 branch-stem loop, which drives strand displacement through a loaded-spring mechanism. This work reveals the physical basis of U2-mediated branch-site recognition and provides a dynamic framework for understanding early spliceosome assembly.

## METHODS

### Model Building

The model of intron-BSL model with the putative three base-pairs-long toehold was built on the cryo-EM structure of human 17S U2 snRNP/SF3b complex (PDB ID 7EVO [14]), considering the BSL structure only. The intron was instead modeled as an A-RNA helix using Nucleic acid builder of AmberTools [24]. The resulting helix was docked onto the BSL by aligning the three bases of the intron complementary to toehold of the U2-BSL. The remaining strand of the A-RNA helix was removed. In the resulting model (**Figure 2A**) two base pairs of the toehold (U37 and G36) formed Watson-Crick base pairs with the intron (A-4 and C-3), i.e. A-4:U37 and C-3:G36. Conversely, one base pair, U-2:A35, was slightly distorted and formed only one hydrogen-bond. Steric clashes between the 5′-end of the intron and U2-BSL were manually resolved by remodeling the intron structure.

**Figure 2:**
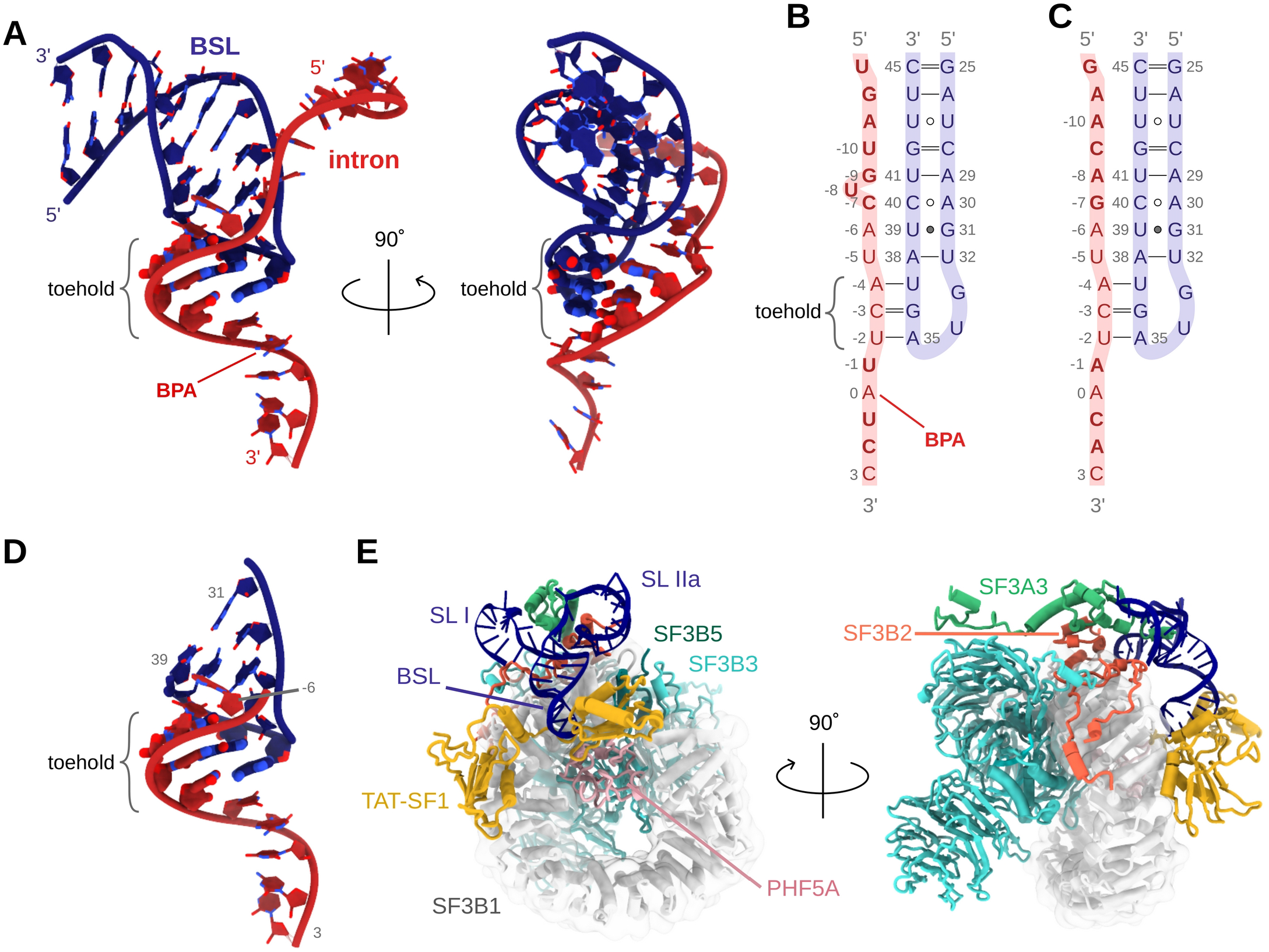
Models systems used for strand displacement simulations. **(A)** Intron-Branch Stem Loop (BSL) model with the intron and the BSL depicted in red and blue ribbons. Base pairs forming the toehold are shown with ticker sticks. Hydrogen atoms are omitted for clarity. Schematic picture of the intron-BSL model with an intron sequence having partial **(B)** and **(C)** full base pair complementarity to the-BSL sequences. Residues differing in **(B)** and **(C)** are highlighted in bold. **(D)** Small intron-BSL construct retaining the only intron positions from −6 to 3. **(E)** 17S U2/SF3b model system. Here the SF3B1, SF3B3, SF3A3, SF3B2 and TAT-SF1 proteins are shown in white, cyan, green, orange and yellow cartoons, respectively.

We built two intron-BSL RNA models considering intron sequences with partial and full complementarity to the U2 snRNA (**Figure 2B,C**). In the first model the intron had the so-called MINX sequence, commonly used in experimental studies. We additionally considered a small model, containing only intron residues from −6 to +3 and BSL residues from 31 to 39 ( **Figure 2D**). This model was created to facilitate the formation of two base pairs in the 5′-end direction of the intron.

Finally, we built a model of the human 17S U2/SF3b system containing only U2 snRNA residues 12-65 (SL I, BSL, and SL II stem-loops), the SF3B1, SF3B3, SF3B5, PHF5A, and TAT-SF1 proteins, and portions of SF3A3 and SF3B2 adjacent to RNA. This model was based on the structure reported in PDB ID 7EVO (**Figure 2E**). Missing protein loops were modeled using RoseTTA [25] and the missing U2 snRNA residues of SL I loop were modeled with the SimRNA webserver [26]. The N- and C-termini of PHF5A were capped with acetyl and methylamine groups (ACE and NME residues in Amber), respectively. One set of the 17S U2/SF3b simulations was run with the TAT-SF1 protein, and another set with TAT-SF1 removed.

### Classical MD simulations

Topologies of the different models were built using the t-leap module of the AmberTools suite. AMBER force fields OL3 [27], [28], [29], [30] and ff14SB [27], [28], [31], [32] were used for RNA and protein, respectively. The molecules were solvated into octahedral boxes of TIP3P water [33] with a minimal distance between the solute and box border of 12 Å for RNA-only systems, and 20 Å for the 17S U2/SF3b complex. Excess KCl salt of 0.15 M concentration was added to all systems; the monovalent ions were described using Li&Merz 12-6 parameters [34]. Zinc ions, coordinated by the PHF5A and SF3A3 proteins in the 17S U2/SF3b complex, were covalently bound to the protein. The ZAFF force field [35] was used for the zinc sites, and the parameters for the zinc site of SF3A3 were derived following the ZAFF protocol available at https://ambermd.org/tutorials/advanced/tutorial20/ZAFF.php.

Overall, the solvated systems contained about 36000-37000 atoms and 328000-390000 atoms for the RNA-only and the 17S U2/SF3b systems, respectively.

To accelerate sampling of the unbiased MD simulations, we also employed two modified variants of the AMBER force field. Namely, we considered hydrogen bond (HB-fix) [36], [37] and stacking-fix (sta-fix) [38] modifications. In the first variant, the short-ranged HB-fix potential adds an energetic barrier (1 kcal/mol) on breakage of the formed hydrogen-bond. Notably, this barrier applies only when the hydrogen bond is spontaneously formed, thus contributing to stabilize its formation without preventing its breakage. The HB-fix potential was applied on the base pairs of the branch helix that were expected to form, including the toehold. In this manner, the conformational equilibrium of the RNA was shifted towards formation of the branch helix without performing a biased simulation. Instead, the sta-fix modification, developed to prevent excessive RNA self-interactions and to enhance sampling, scales down van der Waals interactions between specific RNA atoms. This allows for a more efficient exploration of conformational space. Here, the scaling factor of 0.9 was applied to all RNA nucleotides.

Additionally, to prevent spurious intron circularization, namely sticking of the 5′ and 3′-ends of the intron together, MD simulations of the large intron-BSL construct constructs (**Figure 2A**) were done by applying a wall of 30 Å to end-to-end distance of phosphorus atoms of the intron 5′ and 3′-termini.

Each system was equilibrated following an established protocol [39], which consisted in 11 steps of minimizations and short MD runs with positional restraints decreasing from 5 to 0.5 kcal/mol. This was followed by a 1 ns-long unbiased MD simulation. After equilibration, the production MD simulations were run using the Gromacs program [40] with a 2 fs integration step, keeping the bonds involving hydrogen restrained with the LINCS algorithm. Electrostatics in the periodic boundary field were treated with Particle Mesh Ewald method with a cutoff of 10 Å for short-range interactions. The temperature of 298.16 K was maintained with the v-rescale thermostat [41], while the Parrinello-Rahman barostat [42] was used to control the pressure.

For each system, we ran several replicas. Namely, for the intron-BSL constructs (**Figure 2A**) we ran four replicas with standard (unmodified) force field, and four replicas with HB-fix variant for each intron sequence. Additionally, when using simultaneously the sta-fix and HB-fix force field variants we ran eight replicas with the partially complementary intron sequence and four replicas for the fully complementary intron sequence. This yielded 28 MD simulation trajectories. A summary of the MD simulations performed is reported in **Table S1** of the Supplementary Information. At the beginning of each replica, the velocities from the equilibrated states were randomized to heterogenize the sampling.

All MD simulations of intron-BSL construct were run for at least 500 ns and then, after a visual inspection, either prolonged up to 2 μs or terminated when the trajectories became trapped in states preventing formation of the branch helix. Since most base pairing events, leading to the formation of the branch helix, occurred in the early simulation stages (**Table S2**), we argue that running multiple short MD simulation replicas allows a better sampling of branch helix formation as compared to few longer MD simulations.

MD simulations of the 17S U2/SF3b model were run in two replicas for the complex with the TAT-SF1 protein, and in four replicas for the complex where the TAT-SF1 protein was removed. All replicas were simulated for 2 μs. Helical parameters were calculated using x3-DNA program [43].

### Metadynamics simulations

To estimate the energetics associated to the formation of the branch helix at three base pairs flanking the toehold we performed well-tempered metadynamics simulations [44]. Here, we considered two small RNA-only models, with the intron having either with full or partial base pair complementarity to BSL. These models contained U2-BSL’s nucleotides from U28 to G42 and intron’s nucleotides from C/G-7 to C-3. To increase their stability, the models were capped with one G=C pair at BSL terminus, and by another G=C pair at the toehold, effectively replacing the native U27:U43 and U-2:A35 base pairs of the BSL and intron:BSL duplex, respectively (**Figure S1**).

The well-tempered metadynamics simulations were run using two collective variables (CVs). Namely we applied the εRMSD CV to both BSL and branch helix. εRMSD is a geometrical descriptor capturing the relative positions and orientations of the bases with respect to a reference structure [45]. This CV distinguishes different base-pairing patterns better than the canonical RMSD metric. For this reason, εRMSD is routinely applied to study conformational remodeling of nucleic acids. As a reference structure for εRMSD to BSL, we used the structure deposited in PBD ID 7EVO [14]. Conversely, the εRMSD to the branch helix with partial complementary intron was calculated using a reference structure deposited in PDB ID 7QTT [46]. Finally, the εRMSD to the branch helix, formed with a fully complementary intron sequence, was calculated with respect to a reference canonical A-RNA helix structure built by Nucleic acid builder of AmberTools. Aiming to simulate the initial steps of strand invasion, we only biased the disruption/formation of the first three base pairs beyond toehold. Namely, these are intra-BSL base pairs U32:A28, G31:U39, A30:C40, and branch helix base pairs U-5:A38, A-6:U39, and C/G-7:C40. The εRMSD cutoff in metadynamics simulation was set to 3.2 to cover large areas of the conformational space [47]. To build the bias, we used Gaussian hills with a width of 0.1, deposited every 200 steps on a grid ranging from 0 to 5. The bias factor was set to 10 and τ, controlling the Gaussian hill height, of 70. The simulations were run using Plumed-patched Gromacs [48].

To avoid that the system remained trapped in states distal from the target one, we applied wall restraints to restrict the conformational space sampled during the simulations. Namely, we applied walls restraint on (i) the hydrogen bond distances of the base pairs that were not involved in the BSL-branch helix exchange; (ii) the distance of the two base pairs enclosing the biased bases (A-4:U37 and A29:U41); (iii) on the backbone conformation of the intron strand using pseudo-dihedrals defined over C4’ and P atoms [49]; (iv) a upper wall on the cumulative εRMSD; and (v) a lower wall on (εRMSD_BSL_ – εRMSD_BH_)^2^ – min(εRMSD_BSL_,εRMSD_BH_)^2^ value. The four metadynamics simulations reported here differ in the extent of backbone walls applied. Namely, in the first replica of the partially complementary intron sequence we applied a restraint wall on the backbone of C-7 and A-6 bases, while in the second replica the restraint wall was extended also to the U-5 and A-4 bases. Conversely, in the first replica of a fully complementary intron, we applied analogous restraint walls to G-7, A-6, U-5, and A-4, while in the second replica, the same pseudodiherals were targeted, but they were restrained to a narrower value range. Details are listed in Supplementary Information, Supplementary Methods section. Application of these restraints improved the sampling of BSL-branch helix transitions, but did not completely prevent the sampling of off-pathway states. Thus, we discarded the portions of trajectory corresponding to off-pathway states from the analyses (see Supplementary Methods for details).

Free energies were calculated using reweighting. We considered as belonging to a specific free energy minimum all frames with εRMSD to the reference state lower than 0.7. A εRMSD cutoff of 2.4 was used for analysis. For the partially complementary intron, the free energy was evaluated using a time-averaged bias potential. The averaging was done over the full trajectory except the first 1 µs. For the fully complementary intron, raw (non-averaged) bias files were used due to lower convergence. The errors were estimated using bootstrap analysis.

### Coarse-grained simulations

For coarse-grained simulations we have used the oxRNA model [50], which approximates nucleotides as rigid bodies with two beads – one for the base, and one for the backbone. This model has been successfully used for strand displacement simulations [51], [52]. oxRNA defines interaction potentials for A:U, G:C, and G:U base pairs only. Therefore, we performed these simulations only considering the fully complementary intron sequence. The starting coordinates and topology were generated from an all-atom structure, as shown in **Figure 2A**, using the oxView server [53]. We then ran CG simulations in 100 replicas in which the structures were first relaxed via Monte Carlo and molecular dynamics, followed by a production run for 25×10^6^ steps using a time step of 0.001 in the oxRNA internal time units. This formally corresponds to 750 ns. However, the CG nature of the model makes a direct comparison to real timescales nontrivial, and the actual corresponding physical timescales are typically significantly longer [50], [54]. Therefore, we focused only on the relative timing of individual trajectories in branch helix formation, without attempting to assign physical time scales to CG trajectories.

The resulting trajectories were analyzed using the OAT package [55]. A base pair was considered formed if its hydrogen-bond energy was below −0.2 oxRNA internal units. The BPA was considered to be bulged out if its neighboring base pairs A-1:U34 and C1:G33 were formed, the stacking energy of A-1/C1 pair and of U34/G33 pair was below zero, and the stacking energy of A0 and both its neighboring residues (A-1 and C1) was zero.

## RESULTS

### A slip-stranded conformation of BSL triggers branch helix formation

To monitor the initial steps of the intron strand invasion into the U2-BSL we considered two different models, differing in the extent (partial or full) of the base pair complementarity of the intron to the BSL (**Figure 2B** and **C**). The starting structures accounted only for the initial binding of the toehold. Namely, the U-2,C-3 and U-4 bases of the intron were paired to the A35, G36 and U37 nucleotides of the BSL, respectively. For each model we performed multiple MD simulation replicas of 0.5-2.0 μs, considering three different force-field settings. We used the standard AMABER OL3 force field (FF) [27], [28], [29], [30], a variant of this FF, which stabilized all hydrogen-bonds of the branch-helix base pairs (HB-fix FF variant) [36], [37], and a FF variant in which RNA self-interactions were also weakened (combining HB-fix and sta-fix corrections) [38].

MD simulations, performed with the standard AMBER OL3 FF, revealed the U-2:A35 base pair of the toehold was the least stable among the three toehold base pairs. Indeed U-2:A35 remained fully stable only in 38 % of the MD simulations (3/8 runs), likely due to a geometric constraint of the BSL loop on the A35 backbone conformation. Instead, the two remaining bases G36 and U37 of the toehold formed stable interactions with C-3 and A-4 of the intron, respectively. As expected, a larger stability of all toehold base pairs was instead observed when introducing the HB-fix variant of the FF.

Importantly, 39% of all simulations (11/28) sampled spontaneous toehold extension. Moreover, while with the standard AMBER FF only the U-5:A38 base pair formed in the intron 5′-end direction, the modified FFs aided the formation of two (HB-fix) and four (HB-fix and sta-fix) new base pairs (**Table S2**). Notably, in most of these trajectories (8/11), we observed the formation of a slip-stranded BSL intermediate preceding the toehold extension (**Figures 3** and **4**). The same BSL slip stranded state was also observed in 5 additional simulations, although they did not result in a productive toehold extension. The slip-stranded state featured an intra-molecular register shift in the BSL base pairing, which resulted in a shift by one base of the BSL 3′-strand towards 5′-end. As a result, the A38:U32 base pair of the BSL broke, the U32:U39 pair newly formed, thus liberating A38 for the first step of branch helix extension ( **Figure 3**). This slip-stranded conformation also aided the subsequent BSL displacement step. Here, U39 of the BSL formed a weak intra-BSL base pair with U32, facilitating the binding of U39 to the intron A-6. After formation of the slip-strand state, a propagation of the register shift to the flanking bases was commonly observed. This induced the formation of a second G31:C40 and, occasionally, of a third A30:U41 slip-stranded pair (**Figure S4**). Upon formation of A30:U41 base pair, A29 remained unbound, inducing a destabilization of the C28:G42 base pair, with resulted in C28 bulging. Sometimes, the formation of the slip-stranded states was only short-lived (i.e., ns-long for the U29:U39 pair). Yet, this was sufficient to trigger the formation of the new branch helix base pair.

**Figure 3:**
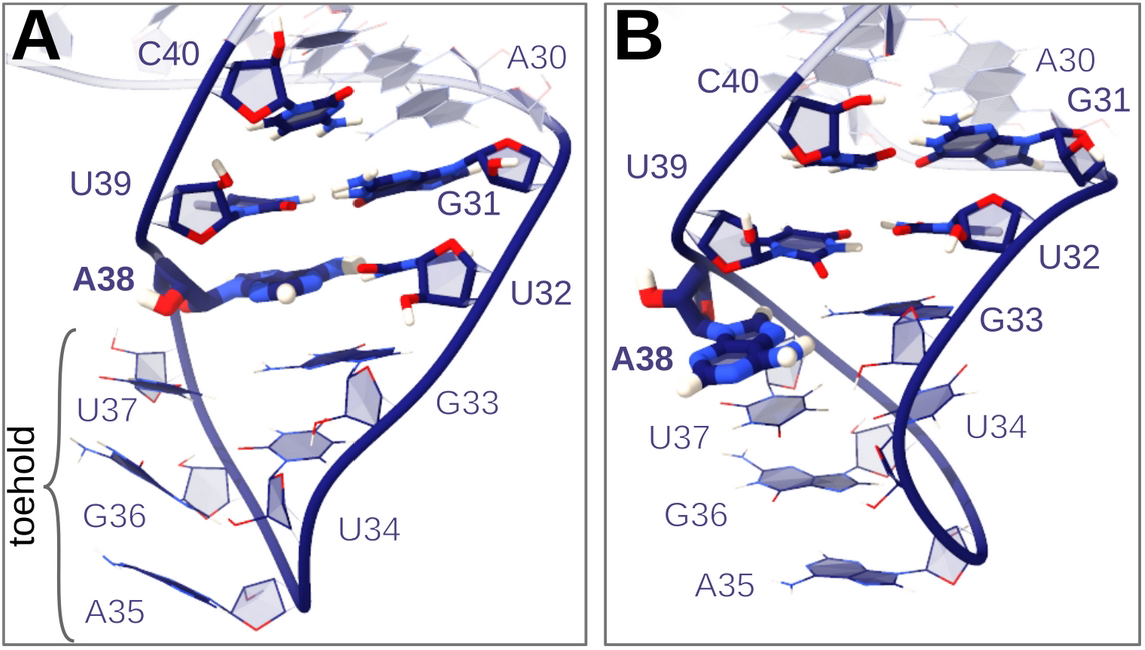
Conformational rearrangement of the BSL. **(A)** The initial conformation and **(B)** the slip-stranded state. Residues involved in the formation of two slip-stranded base pairs (U32:U39 and G31:C40) are shown in sticks, the rest in lines. Intron, bound to the toehold residues A35-U37, is not shown for clarity.

Conversely, in few simulation replicas, where the initial slip-stranded event was not sampled, the formation of first two base pairs (U-5:A38 and A-6:U39) of the branch helix occurred immediately after the opening of the A38:U32 and U39:G31 intra-BSL base pair. Occasionally, the BSL bases transitioned from intra-BSL to BSL-intron base pairing patterns by forming temporary interactions with both the BSL and intron strands. This mechanism also applied to the formation of the third (C/G-7:C40) and fourth (A-8:U41) base pairs of the branch helix.

Conversely, no extension of the branch helix was observed at the other extremity of the toehold (intron 3′-direction). There, only the A/U-1:U34 base pair transiently formed in two simulation replicas.

To further examine the occurrence of the slip-stranded pathway, we performed MD simulations on a smaller intron-BSL construct. This contained only two bases beyond the toehold (i.e., intron residues from −6 to +3 and BSL residues from 31 to 39, **Figure 2D**). The reduced size of the model enabled the spontaneous formation of two new branch helix’s base pairs even without introducing any FF modifications (**Figure S5**). Importantly, the initial slip-stranded state of BSL preceded the formation of the first branch helix base pairing, consistently with simulations of the full-size intron-BSL model.

### Early steps of branch helix formation are energetically favorable

To characterize the energetics of branch helix formation, we performed two metadynamics simulations on the reduced BSL-intron model, considering intron sequences partially and fully complementary to the BSL (**Figure 5A**). Each metadynamics simulation of the partially complementary intron sequence sampled 13 transitions of BSL to the branch helix, (Supplementary Methods), which resulted to be energetically favorable (the calculated free energy differences between the starting BSL and final branch helix states (**ΔG**_**BH-BSL**_) were −4.6**±**1.3 and −3.4**±**1.9 kcal/mol in the two replicas). This process became even more energetically favorable when considering the fully complementary sequence (**ΔG**_**BH-BSL**_= −9.3**±**4.8 and −5.0**±**3.1 in the two replicas). However, with this model the BSL to branch helix transition was sampled only 4-5 times per simulation. Hence, due to their more limited sampling of these transitions, these metadynamics simulations are affected by larger errors. As such, although these simulations can only provide qualitative information, they still suggest that, irrespectively of the intron sequence, the formation of the first branch helix base pairs is thermodynamically more favorable than the intra-BSL base pairing.

**Figure 4:**
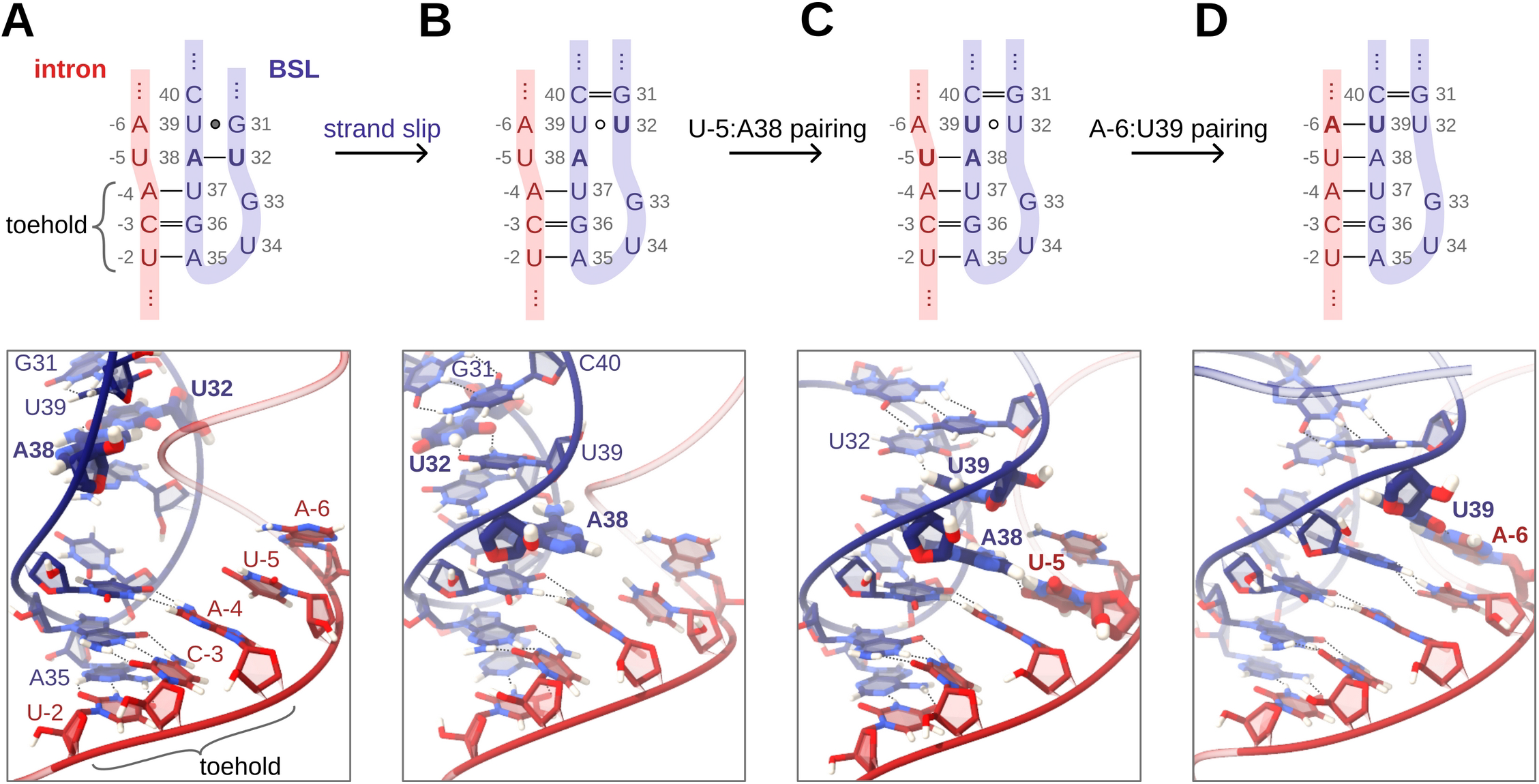
Early steps of branch helix formation via the slip-stranded path. Schematic representations (top panel), and snapshots (bottom panels) of the intermediates observed in all-atom simulations. Residues involved in the toehold and BSL (blue):intron (red) remodeling steps are shown in sticks, the remaining residues are not shown. **(A)** A three-base-pair toehold, based on the base pairing interactions between A-4:U37, C-3:G36 and U-2:A35 of the intron and BSL, respectively, is initially present. The BSL contains the U32:A38 and G31:U39 base pairs. **(B)** BSL transits to the slip-stranded conformation with formation of U32:U39 and G31:C40 base pairs, while A38 remains unpaired and protrudes towards the intron. **(C)** A38 pairs with U-5. **(D)** U39 pairs with A-6 further extending the branch helix.

**Figure 5:**
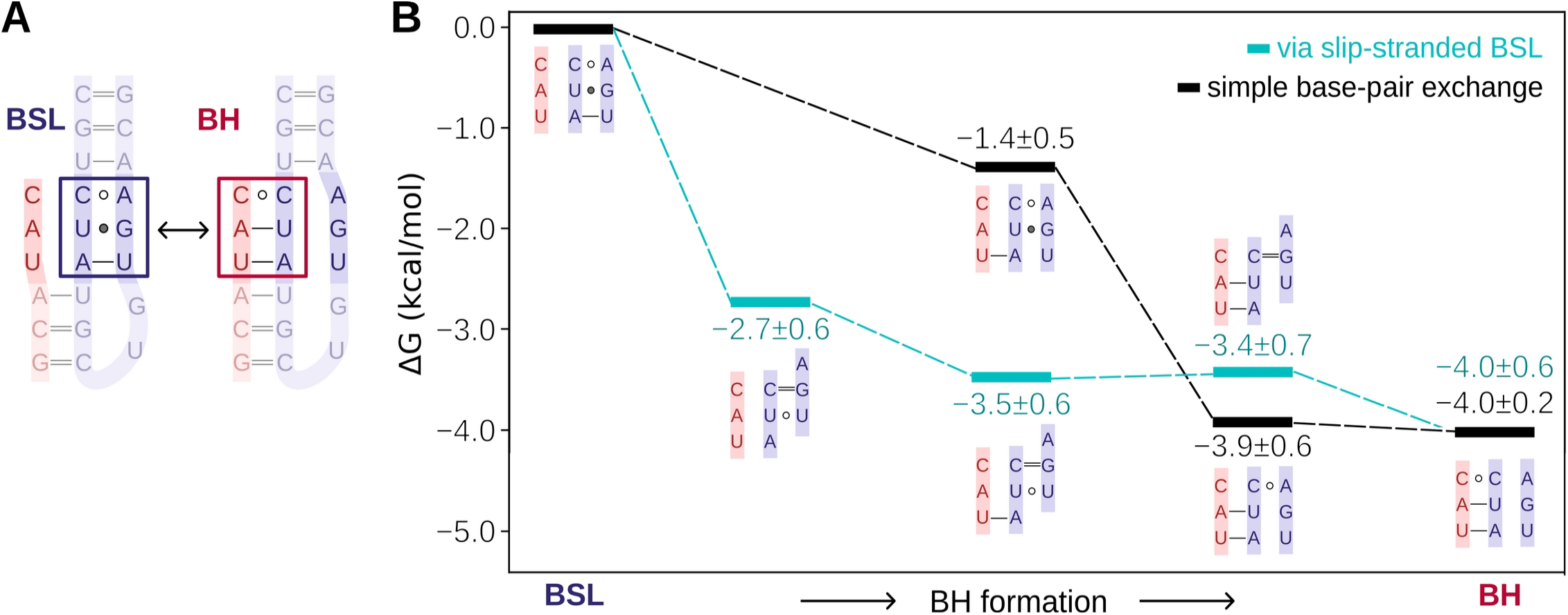
Energetics of the first three steps of toehold extension. **(A)** Model system used in metadynamics simulations featuring an intron (red) with partially complementary sequence to the BSL (blue). The boxes mark the bases downstream the toehold that were biased in metadynamics simulations. **(B)** Scheme of the relative free energy (kcal/mol) of the two competitive pathways (slip-stranded (cyan lines) and base-pair exchange (black lines)) leading to the formation of the branch helix (BH). The scheme shows only the residues involved in interaction remodeling. Free energies (kcal/mol) are averaged over two metadynamics replicas (see **Figure S6** for individual runs).

Spontaneous formation of the slip-stranded state was also sampled in metadynamics simulations, despite not being explicitly enforced by the collective variables used. Notably, the first and the second steps of the slip-stranded pathway (register shift of the U-5:A38 pairing) are energetically more favorable than the starting BSL conformation (**Figure 5**, DG=-2.7**±**0.6 and −3.5**±**0.6, respectively), with the first step being the most energetically favorable. Conversely, the following steps of toehold extension are, within the error of our calculations, isoenergetic. An alternative path, which avoided the formation of slip-stranded intermediates, was also sampled. As discussed above, this consisted in an exchange of intra-BSL base pairs in favor of intron:BSL base pairs. In this base pair exchange path, the formation of first (U37:A-4) and the second (A38:U-5) base pair was thermodynamically favorable (−1.4**±**0.5 and −3.9**±**0.6, respectively). Importantly, the free barriers for the conversion between states were small (about 1-2 kcal/mol) in both the slip stranded or base pair exchange paths, suggesting that all states were thermally accessible. Thus, although the slip-stranded pathway appears to be slightly more favorable, both the slip stranded and the base exchange pathways seem kinetically and thermodynamically viable at room temperature.

### TAT-SF1 prevents formation of the slip-stranded state of BSL in the 17S U2/SF3b complex

We next assessed if the slip-stranded BSL conformation could form even within the SF3b protein environment. Specifically, to monitor the impact of TAT-SF1 on BSL stability and its role in preventing the formation of the branch helix, we simulated the 17S U2/SF3b particle (**Figure 2E**) either in presence or absence of TAT-SF1. The model without TAT-SF1 represents a hypothetical state formed after TAT-SF1 dissociation aided by PRP5, but in which the intron is not yet bound to SF3b.

During multi-replica 2 µs-long MD simulations the U2/SF3b core was structurally stable, while the TAT-SF1 protein, BSL, and SL I stem loop of U2 snRNA experienced large fluctuations, consistently with weak electron densities observed in cryo-EM experiments [14], [56] (**Figure S7**). While the presence of TAT-SF1 reduced fluctuations of the BSL, its removal triggered a BSL restructuring (**Figure 6**). Namely, in two MD simulation replicas, where TAT-SF1 was absent, sampled the formation of the strand-slipped BSL. Conversely in the two other replicas, we observed (i) a marked restructuring of the BSL via a complete change of the base pairing pattern and (ii) SL I relocation towards BSL, causing a rupture of G25:C45 and A26:U44 base pairs while the region adjacent to BSL loop was stable. As such, MD simulations importantly reveal that the BSL slip-stranded state can also form within the 17S U2/SF3b complex.

**Figure 6:**
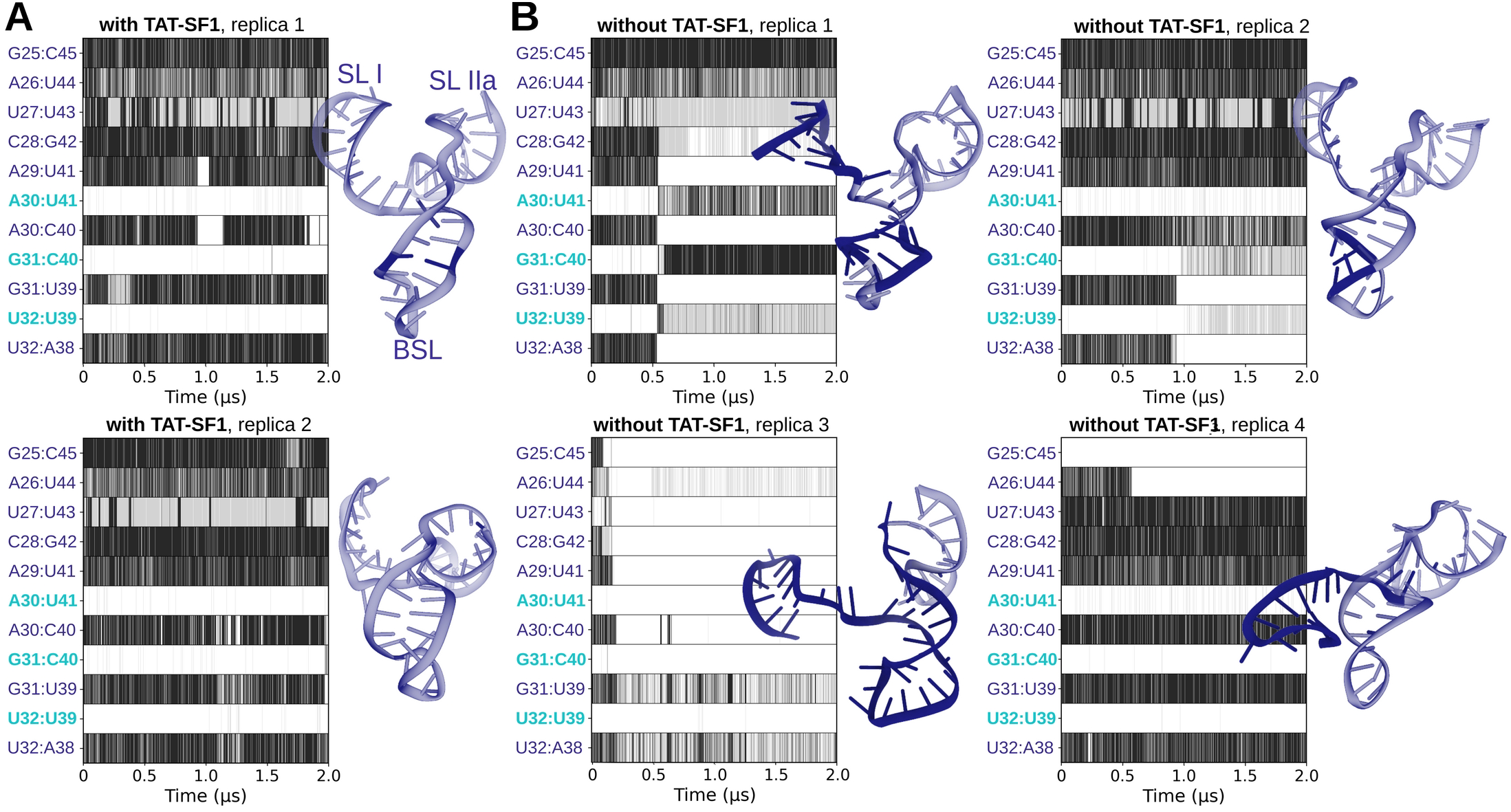
Dynamics of the branch stem loop (BSL) within the 17S U2/SF3b complex. Molecular Dynamics simulations of the 17S U2/SF3b complex were done with **(A)**, and without the TAT-SF1 protein **(B)**. Graphs show the evolution of base pairing formation versus simulations time (μs) in different models and replicas. The base pairs labeled in cyan participate the strand-slipped conformation. Black, gray and white colors indicate that the base pair is fully formed, weakly-bound (suboptimal pairing geometry, lower number of hydrogen-bonds), and absent, respectively during the MD simulation trajectory. The U2 snRNA structures sampled in the last snapshot of each trajectory is shown on the sides of the trajectory in blue color with restructured bases highlighted in a darker tone.

The spontaneous and readily accessible BSL rearrangements observed in the absence of TAT-SF1 suggest that this protein might keep BSL in a structurally tensed state. Indeed, an analysis of the helical parameters revealed that TAT-SF1 stabilized a different BSL conformation, overtwisted and characterized by a larger helical rise and radius. This contrasts notably with the BSL, isolated or within 17S U2 snRNP that lacks the TAT-SF1 protein (**Figure 7**).

**Figure 7:**
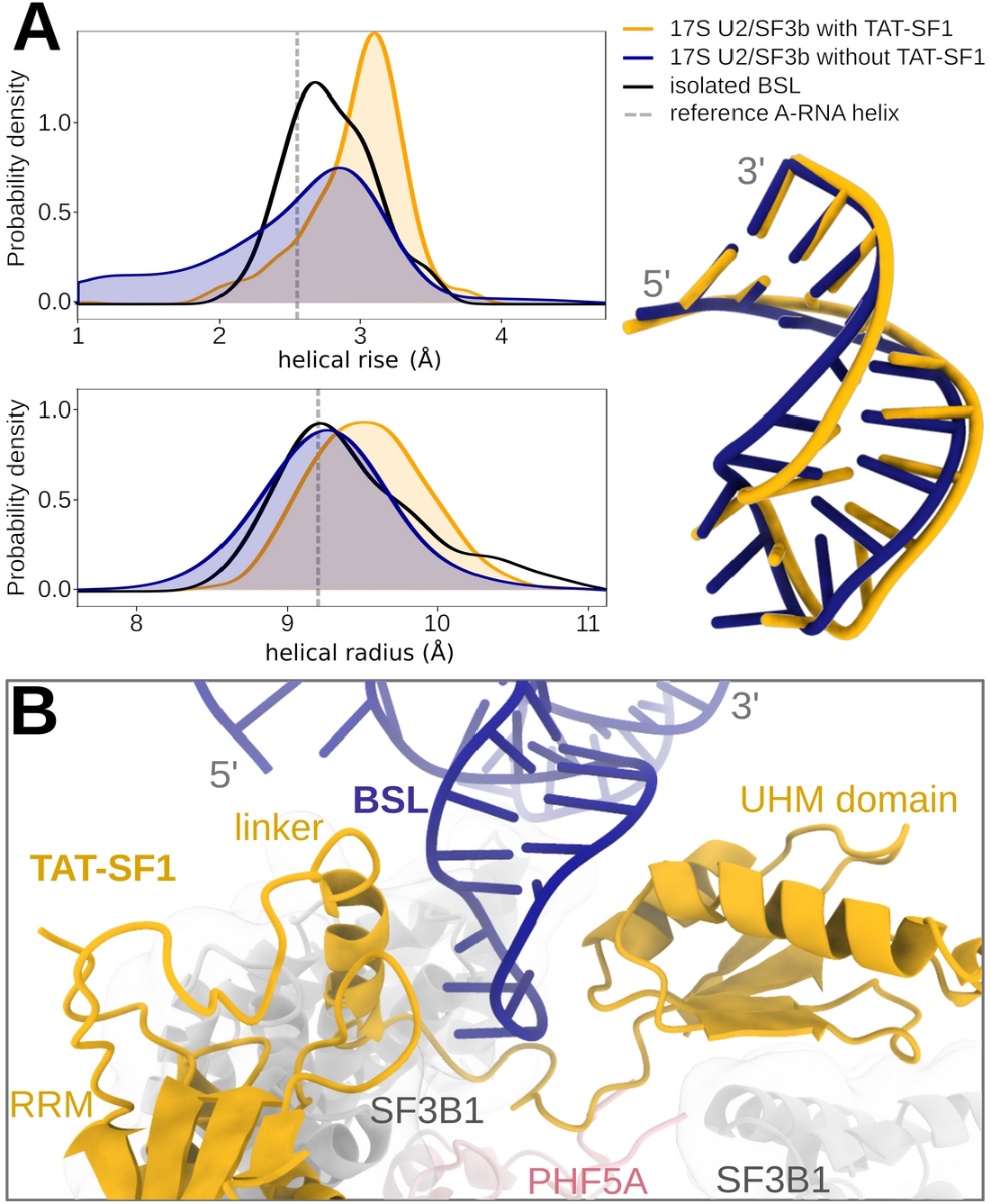
BSL conformation in the presence and absence of TAT-SF1 protein. **(A)** Helical parameters of the BSL stem. Left panel shows distribution of helical rise values, defined as the distance between the planes of the consecutive base pairs and of the helical radius values, defined as the distance between helical axis and phosphorus atoms. The plots contain data from two merged trajectories of 17S U2/SF3b with TAT-SF1 (yellow), four merged trajectories of 17S U2/SF3b without TAT-SF1 (dark blue). For a reference, data of a 250 ns-long trajectory of isolated BSL (black) are also shown. Reference values for A-RNA helix were taken from Ref. [43]. Right panel shows the corresponding average BSL structures with (yellow) and without (blue) the TAT-SF1 protein. Structures are aligned with respect to the two terminal base pairs. **(B)** Structure of the 17S U2/SF3b with TAT-SF1(yellow) masking the BSL (blue). See also **Figure 2E**.

### Coarse-grained simulation of complete branch helix formation

We finally attempted to simulate the formation of the full branch helix. Due to the complexity and the time scale required for this substantial conformational remodeling we accelerated the sampling by performing CG simulations with comes at the cost of losing atomic-level details. We choose the oxRNA CG model, since this was already successfully applied to study strand invasion [51], [52]. However, since this model account only for canonical and G:U wobble base pairs, we simulated only the intron sequence having fully complementary to the BSL. GC simulations were started from the same toehold-bound structure used in the all-atom MD simulations (**Figure 8**). By performing 100 CG simulation replicas we observed in vast majority of trajectories (83 runs) the complete formation of the branch helix. In 10 trajectories the branch helix formed only partially, and in 7 trajectories the toehold was lost and the two strands dissociated.

**Figure 8:**
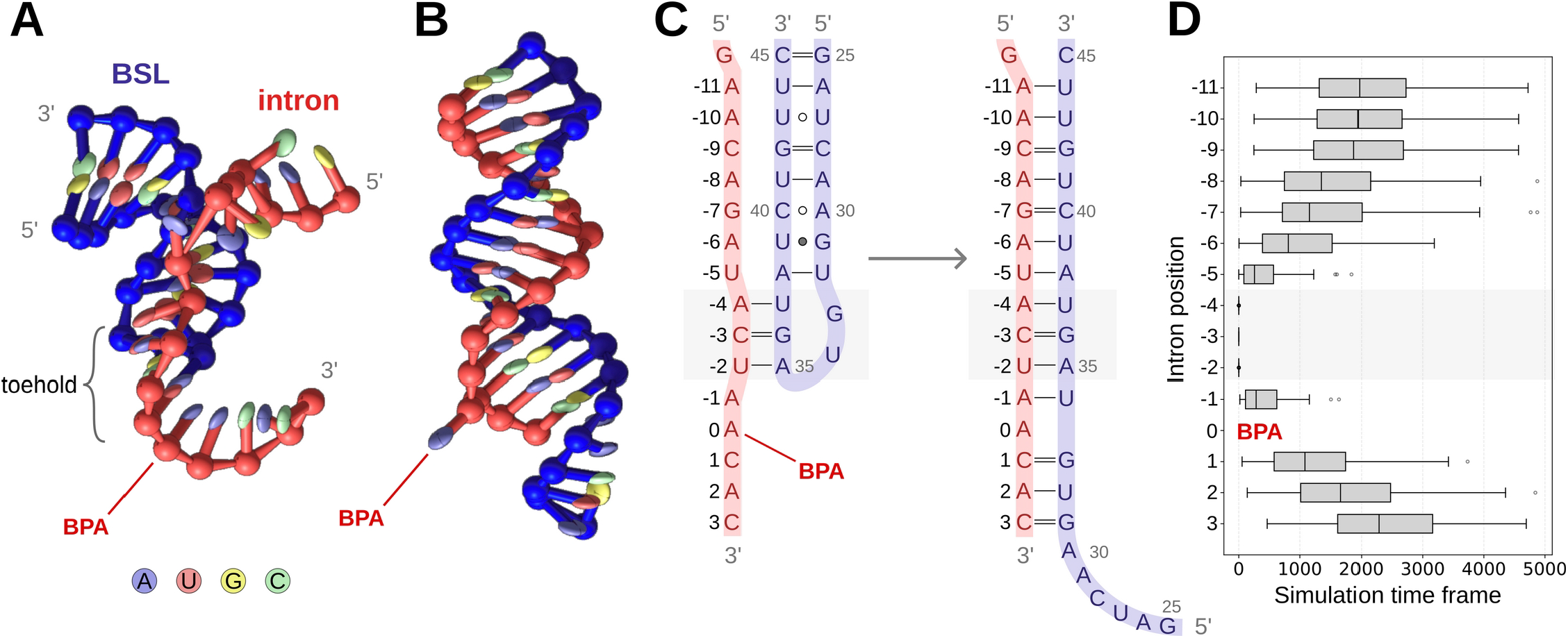
Branch helix formation in coarse-grained (CG) simulations. Starting **(A)** and final **(B)** structure. In the selected snapshot of the final structure the BPA is bulged out. **(C)** Schematic representation of **(A)** and **(B)**. The toehold area is marked by a gray box. **(D)** Statistics on the branch helix base pair formation vs. simulation time frames. The average is collected over 100 CG trajectories.

Consistently with all-atom simulations, the formation of the branch helix proceeded in a step-wise manner. Only the terminal portion of the BSL (i.e., involving intron nucleotides from position −9 to −11) formed concertedly with complete BSL stem unwinding (**Figure 8D**). The growth of the branch helix started from the bases flaking the toehold and proceeded linearly in both directions at a similar speed (**Figure 8D**). The slip-stranded state was not sampled in CG simulations most likely due to inability of the oxRNA model to capture non-canonical base-pairs.

Additionally, in 30% of the trajectories the formation of a transient BPA bulge-out events was observed. During the growth of the branch helix the BPA remained intrahelical and occasionally paired with U34 or G33, being budged out only after complete formation of the branch helix (**Figure S8**).

## DISCUSSION

RNA conformational remodeling plays crucial roles in many pivotal biological processes, including translation regulation, protein synthesis, and pre-mRNA splicing [57], [58]. During remodeling, RNA conformations typically adopt two mutually exclusive sets of base pairing interactions. The transitioning from one conformation to another requires breaking one set of base pairing interactions in favor of the new set. While RNA helices can be unwound by helicases in an ATP-dependent manner [59], [60], [61], RNA restructuring can also occur through a non-ATP-driven strand displacement mechanism [2]. In this process, a spontaneous dissociation of an RNA helix, which requires overcoming a high free energy barrier, is unlikely to serve as the primary mechanism for RNA remodeling. An alternative stepwise process, based on a progressive exchange of interactions, is more energetically viable and thus most likely to take place. In this category falls the strand displacement mechanism in which a single RNA strand invades a duplex, displacing one of the duplex strands to hybridize with the other.

While RNA strand displacement is well characterized *in vitro* and used for nanotechnological applications [3], [62], its characterization in cellular content lags. Among biological systems suggested exerting their biological function via strand displacement is the CRISPR-Cas system, where the spacer region of the guide RNA leads the complex to the target genome region and disrupts the DNA:DNA duplex to form an RNA:DNA duplex, a process known as R-loop propagation [63], [64]. RNA rearrangement also occurs during co-transcriptional folding, as the nascent RNA begins to fold while being transcribed. Since the initial folding pattern of the RNA transcript does not necessarily represent the lowest-free energy conformation of the entirely synthesized sequence, the RNA can rearrange its secondary structure trough a strand displacement as the transcription proceeds [65]. Some of the dramatic changes in riboswitch conformation in response to ligand binding were also suggested to occur via a strand displacement process [66]. Some non-coding RNAs were also hypothesized to engage with their targets via strand displacement [67].

Among biological systems proposed to exploit strand displacement, the spliceosome assembly presents a particularly intriguing case. Cryo-EM studies suggest that U2 snRNA recognizes the intronic branch sequence (BS) through a strand invasion process [15]. This mechanism, however, poses a puzzle: how can U2 snRNA spontaneously invade and replace the branch-stem loop (BSL) pairing despite the limited complementarity of typical intron sequences?

Here, we provide a mechanistic answer to this question by revealing the physical basis of strand displacement during U2-mediated branch-site recognition. Using all-atom and coarse-grained molecular dynamics simulations, we show that strand invasion can proceed spontaneously once the toehold region is engaged and the TAT-SF1 protein dissociates. The key discovery is that TAT-SF1 maintains the BSL in a supercoiled, high-energy conformation, acting as a molecular latch that holds U2 snRNA in a “loaded-spring” state. Our modeling supports the scenario where TAT-SF1 release enables relaxation of this strained BSL conformation, unleashing the stored conformational (twisting) energy and causing strand invasion. We also show that the latter can proceed via two pathways. The first pathway (slip stranded path) involves a strand slippage, where the resulting BSL state features a register-shifted base-pairing pattern (**Figures 3** and **4**). This slip-stranded intermediate of the BSL facilitates the readout of the intron at position U/Y-5, which pairs with A38 of BSL. Namely, the BSL remodeling into a slip-stranded state liberates A38 for intron selection and binding to U2 snRNA. In principle, the BSL might adopt the slip-stranded state after the intron binds to the three-base pair toehold of the BSL (**Figure 9**), or before intron binding, thus creating a four-base pair-long toehold. Such small-scale RNA structural switching, involving non-canonical base pairs and/or loop-adjacent areas, occurs spontaneously on μs timescale [68], [69], [70]. The relevance of the here observed strand slippage mechanism is supported by its recurrence in other biological processes such as gene expression control [71] or guanine quadruplex folding [72], [73], [74]. As an alternative, a more conventional base-exchange pathway, can also take place. In this path, an intra-BSL base pairs interactions are progressively lost in favor of inter BSL-Intron base pairs formation. The events triggering the growing of the branch helix are thermodynamically favorable in both the slip-stranded and base-exchange pathways (**Figure 5**). Albeit the formation of the slip-stranded states appears to be slightly more energetically favorable than the base-exchange pathway, both pathways are viable even when the intron has limited base pair complementarity to the BSL. Finally, since BSL stem distant from the toehold-forming region is characterized by a high base pairing complementarity, and it is therefore quite thermodynamically stable, one can assume that the formation of the final portion of the branch helix proceeds at lower pace.

**Figure 9:**
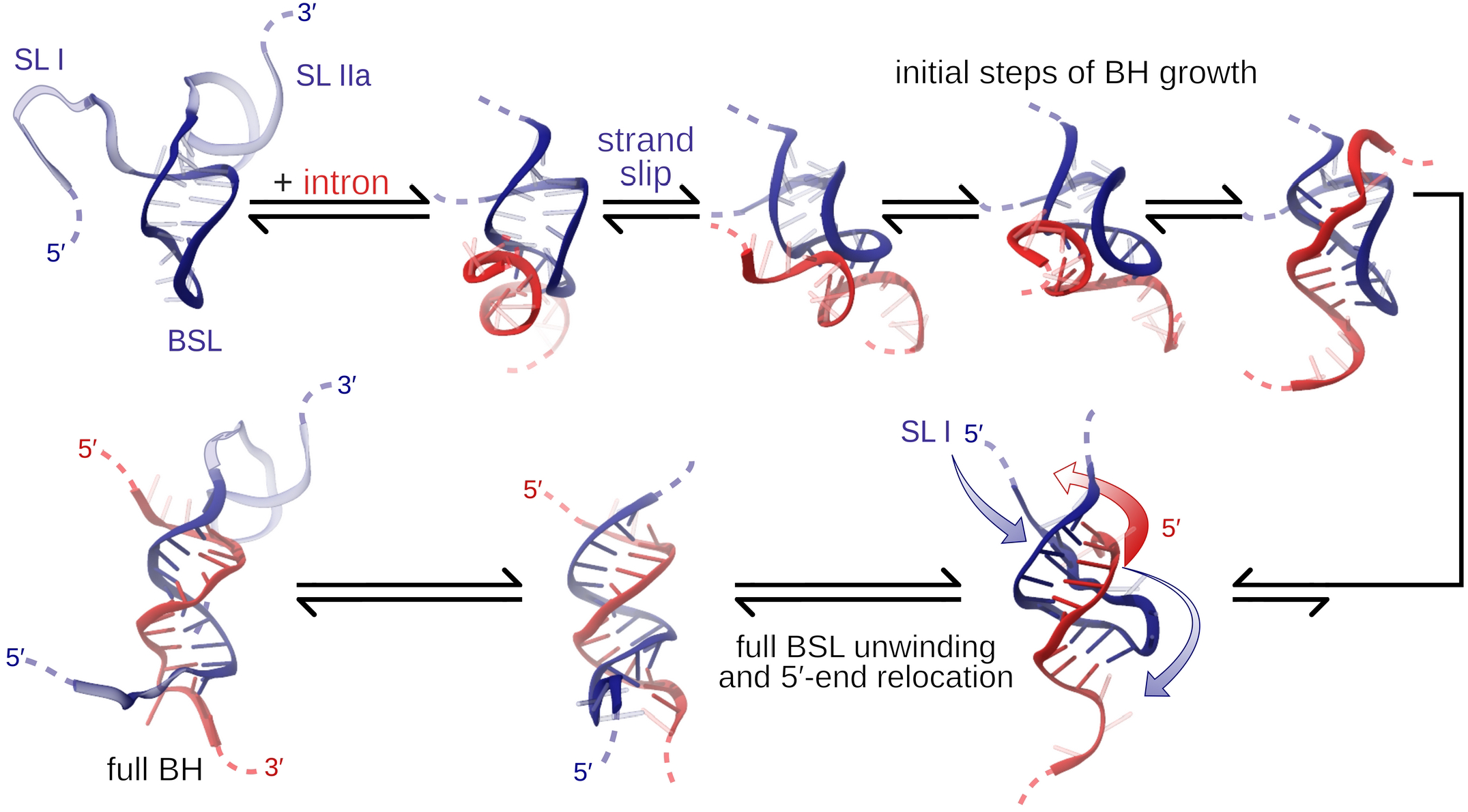
Proposed mechanism of intron:U2 hybridization though slip-stranded pathway. In the first stage, the BS quickly binds to BSL and forms a few new base pairs with the aid of the slip-stranded BSL state (the first line). The rest of the BH extension occurs more slowly, requiring also a swap of the intron and U2 snRNA 5′-ends (the second line, indicated by arrows). Namely, the SL I has to unwind and pass below the intron, while the intron has to pass in the opposite direction. This exchange is restricted by the presence of SL II and SFB1 in the full U2/SF3b system (**Figure S9**).

Our simulations also reveal that strand invasion can proceed bidirectionally, suggesting a dynamic interplay of base-pair exchange along both 5′ and 3′ directions of the branch helix. The unidirectional strand displacement observed by cryo-EM might be thus due to the constraints imposed by the Spliceostatin A, that occupies the BPA binding pocket, therefore acting as an obstacle preventing the bidirectional strand invasion.

Notably, to fully hybridize with the intron, the BSL loop might need to temporarily detach from SF3B1 surface. Shortly, the 5′-end of U2 snRNA, containing SL I, must unwind and pass between BSL 3′-end and SF3B1, while the intron, in turn, must undergo the opposite movement to accommodate in the narrow space between the U2 snRNA and SF3B1 (**Figure 9**). This non-trivial rearrangement is expected to further slow-down the final steps of branch helix formation. However, since BSL and SL I are very flexible, it is possible that, after TAT-SF1 removal, they may dissociate from SF3B1 to allow for the BSL/intron strand exchange before the newly formed branch helix docks back onto the SF3B1.

The mechanistic picture collectively provided by our simulations also aligns with the kinetic proofreading model. The proofreading, underlying BS selection, has been suggested to occur via a kinetic competition between intron binding and U2 snRNA large-scale restructuring. This may either lead to progression of the splicing cycle or to the formation of an inactive state, which arrests the progression [56]. The competition is triggered by TAT-SF1 removal, which upon its dissociation, releases BSL’s structural tension, leading to BSL restructuring. Thus, the BSL conformational switch observed here enables a quick readout of the 5′-segment of the consensus BS sequence, promoting the binding of suitable introns. Otherwise, the formation of an inactive state will take place, and the intron will be rejected [56].

Overall, we reveal that the intrinsic flexibility of the BSL, owing to non-canonical base pairing of the stem, is crucial for its function. BSL plasticity indeed facilitates its conformational remodeling. This mechanism is most likely shared by other biological systems that leverage intrinsic RNA flexibility to enhance the strand invasion. Indeed, the presence of non-canonical, flexible base-pairs was recently demonstrated to enhance strand displacement also in riboswitches [65], [75], [76].

Finally, this work underscores the power of molecular simulations to complement cryo-EM structural data and capture transitions, corresponding to short living intermediates that are difficult to access in purified complexes. By revealing the dynamic and energetic underpinnings of U2 snRNA strand invasion, our study shows the loaded-spring mechanism as critical for branch site recognition, and suggests that it may represent a more general paradigm for understanding how RNA elements harness internal strain to drive conformational remodeling in complex RNP assemblies.

## ACKNOWLEDGMENTS

We thank Wojciech P. Galej for insightful discussions of the studied mechanism and Michael Matthies for his dedicated guidance on coarse-grained simulation setup. A.M. thanks the Italian Association for Cancer Research (project AIRC IG 24514). We further acknowledge the CINECA award under the ISCRA initiative, for the availability of high performance computing resources and support.

## SUPPORTING INFORMATION

Supporting Information PDF file contains Supplementary Methods (details on metadynamics simulations), Supplementary Tables S1-S2 (simulation list and a list of restructuring events), and Supplementary Figures S1-S9 (structures from metadynamics simulations, convergence analysis, detail on the slip-stranded state, analysis of branch helix growth in the simulations of the small system, free energy analysis for individual replicas, RMSF analysis for the U2/SF3b complex, analysis on BPA bulge-out events in coarse-grained simulations, and a scheme of putative RNA restructuring in the U2/SF3b complex).

## DATA AVAILABILITY

All simulation input files, derived parameter files, reference structures and analysis scripts are available in the SI, at https://github.com/ppokor/U2_BH, and at Zenodo database (DOI 10.5281/zenodo.15775838) following the FAIR principles [77].

## Notes

https://github.com/ppokor/U2_BH

https://doi.org/10.5281/zenodo.15775838

### Competing Interest Statement

The authors have declared no competing interest.

