## Supplementary Information PDF for "U2 snRNP recognizes the branch site through a loaded-spring strand-invasion mechanism"

### Supplementary Methods - Metadynamics simulations

Below we list the wall restrains used in the metadynamics simulations, see also Figure S1 and the plumed input files. Notably, test metadynamics simulations performed with additional walls did not improve the sampling of BSL-BH formation events, but induced spurious restructuring in the non-biased segment of the BSL loop instead.

**Metadynamics simulations with intron having partial complementarity to BSL, replica 1:**

1. upper wall with harmonic restraints on X-H…Y H-bond (hydrogen-acceptor distance) for all H-bonds of the four G=C pairs closing the construct (see the Main text for the scheme of the construct): distance of 0.22 nm, force constant of 150 kj/mol
2. upper wall with harmonic distance restraint on center-of-mass of the A-U pairs adjacent to the biased nucleotides: distance of 2.0 nm, force constant of 150 kj/mol
3. pseudodihedral of $C_{-7}$(C4')-$A_{-6}$(P)-$A_{-6}$(C4')-$U_{-5}$(P): lower wall with harmonic restraint on sin(pseudodihedral): sin of 0.0, force constant of 150 kj/mol
4. upper wall with harmonic restraint on cumulative εRMSD for both target regions: sum(εRMSD) of 4.0, force constant of 300 kj/mol
5. lower wall with harmonic restraint on $(\varepsilon RMSD_{BSL} - \varepsilon RMSD_{BH})^2 - \min(\varepsilon RMSD_{BSL},\varepsilon RMSD_{BH})^2$, value of -2.0 force constant of 300 kj/mol. This setting prevents sampling of regions where both εRMSD values are large.

**Metadynamics simulations with intron having partial complementarity to BSL, replica 2:**

1., 2., 4., 5., as defined above

3. set of walls with harmonic restraint on pseudodihedrals, all with a force constant of 150 kj/mol:

   $C_{-7}$(C4')-$A_{-6}$(P)-$A_{-6}$(C4')-$U_{-5}$(P): lower wall on sin(pseudodihedral): sin of 0.0

   $A_{-6}$(P)-$A_{-6}$(C4')-$U_{-5}$(P)-$U_{-5}$(C4'): upper wall on sin(pseudodihedral): sin of 0.0

   $A_{-6}$(C4')-$U_{-5}$(P)-$U_{-5}$(C4')-$A_{-4}$(P): lower wall on sin(pseudodihedral): sin of 0.0

   $U_{-5}$(P)-$U_{-5}$(C4')-$A_{-4}$(P)-$A_{-4}$(C4'): upper wall on sin(pseudodihedral): sin of 0.0

**Metadynamics simulations with intron having fullcomplementarity to BSL, replica 1:**

1., 2., 4., 5., as defined above

3. set of walls with harmonic restraint on pseudodihedrals, all with a force constant of 300 kj/mol:

   $G_{-7}$(C4')-$A_{-6}$(P)-$A_{-6}$(C4')-$U_{-5}$(P): lower wall on sin(pseudodihedral): sin of 0.0; upper wall on sin(pseudodihedral): sin of 0.8

   $A_{-6}$(P)-$A_{-6}$(C4')-$U_{-5}$(P)-$U_{-5}$(C4'): upper wall on sin(pseudodihedral): sin of 0.0

$A_{-6}$(C4’)-$U_{-5}$(P)-$U_{-5}$(C4’)-$A_{-4}$(P): lower wall on sin(pseudodihedral): sin of 0.0; upper wall on sin(pseudodihedral): sin of 0.8

$U_{-5}$(P)-$U_{-5}$(C4’)-$A_{-4}$(P)-$A_{-4}$(C4’): upper wall on sin(pseudodihedral): sin of 0.0

$U_{-5}$(C4’)-$A_{-4}$(P)-$A_{-4}$(C4’)-$C_{-3}$(P): lower wall on sin(pseudodihedral): sin of -0.2; upper wall on sin(pseudodihedral): sin of 0.8

**Metadynamics simulations with intron having fullcomplementarity to BSL, replica 2:**

1., 2., 4., 5., as defined above

3. set of walls with harmonic restraint on pseudodihedrals, all with a force constant of 300 kj/mol:

$G_{-7}$(C4’)-$A_{-6}$(P)-$A_{-6}$(C4’)-$U_{-5}$(P): lower wall on sin(pseudodihedral): sin of 0.0; upper wall on sin(pseudodihedral): sin of 0.8

$A_{-6}$(P)-$A_{-6}$(C4’)-$U_{-5}$(P)-$U_{-5}$(C4’): upper wall on sin(pseudodihedral): sin of 0.0

$A_{-6}$(C4’)-$U_{-5}$(P)-$U_{-5}$(C4’)-$A_{-4}$(P): lower wall on sin(pseudodihedral): sin of 0.0; upper wall on sin(pseudodihedral): sin of 0.8

$U_{-5}$(P)-$U_{-5}$(C4’)-$A_{-4}$(P)-$A_{-4}$(C4’): upper wall on sin(pseudodihedral): sin of -0.2; lower wall on sin(pseudodihedral): sin of -0.2

$U_{-5}$(C4’)-$A_{-4}$(P)-$A_{-4}$(C4’)-$C_{-3}$(P): lower wall on sin(pseudodihedral): sin of -0.2; upper wall on sin(pseudodihedral): sin of 0.8

### Supplementary Tables

**Table S1:** List of molecular dynamics simulations performed.

| Force field modifications | Number of replicas | Range of replica lengths (μs) |
|---|---|---|
| **Intron-BSL RNA construct, intron with partial complementarity** | | |
| - | 4 | 0.5-1.0 |
| HBfix | 4 | 1.0-1.5 |
| HBfix, stafix | 8 | 0.5-1.7 |
| **Intron-BSL RNA construct, intron with full complementarity** | | |
| - | 4 | 0.5-1.0 |
| HBfix | 4 | 0.5-1.0 |
| HBfix, stafix | 4 | 0.5-1.0 |
| **Small intron-BSL RNA construct** | | |
| - | 4 | 1.0 |
| HBfix | 1 | 1.0 |
| **U2 17S construct** | | |
| - | 2 | 2.0 |
| **U2 17S construct without TAT-SF1** | | |
| - | 4 | 2.0 |
| **Metadynamics, intron with partial complementarity** |  |  |
| - | 2 | 3.0-6.0 |
| **Metadynamics, intron with full complementarity** |  |  |
| - | 2 | 1.0-5.2 |
| **intron-BSL RNA construct, intron with full complementarity** | | |
| coarse grained | 100 | $25x10^6$ steps |

**Table S2:** Restructuring events towards branch helix formation in the simulations of the complete RNA model performed with different Amber force field variants. Simulations which did not sample any new branch helix base pair are not listed.

| Simulation force field | Length (μs) | 1st BP formed (ns) | 2nd BP formed (ns) | 3rd BP formed (ns) | 4th BP formed (ns) |
|---|---|---|---|---|---|
| **Intron-BSL RNA construct, intron with partial complementarity to BSL** | | | | | |
| standard | 1.0 | 393 | | | |
| HBfix | 1.5 | 174 | | | |
| HBfix | 1.5 | 136 | | | |
| HBfix, stafix | 1.7 | 97 | 102 | 128 | |
| HBfix, stafix | 0.5 | 20 | | 125 | |
| HBfix, stafix | 0.5 | | 379 | | |
| HBfix, stafix (m) | 0.5 | 149 | 149 | | |
| **Intron-BSL RNA construct, intron with full complementarity to BSL** | | | | | |
| standard | 1.0 | 100 | | | |
| HBfix | 1.0 | 341 | 395 | | |
| HBfix, stafix | 1.0 | 22 | 209 | 14 | 249 |
| HBfix, stafix | 1.0 | 352 | 352 | | |

### Supplementary Figures

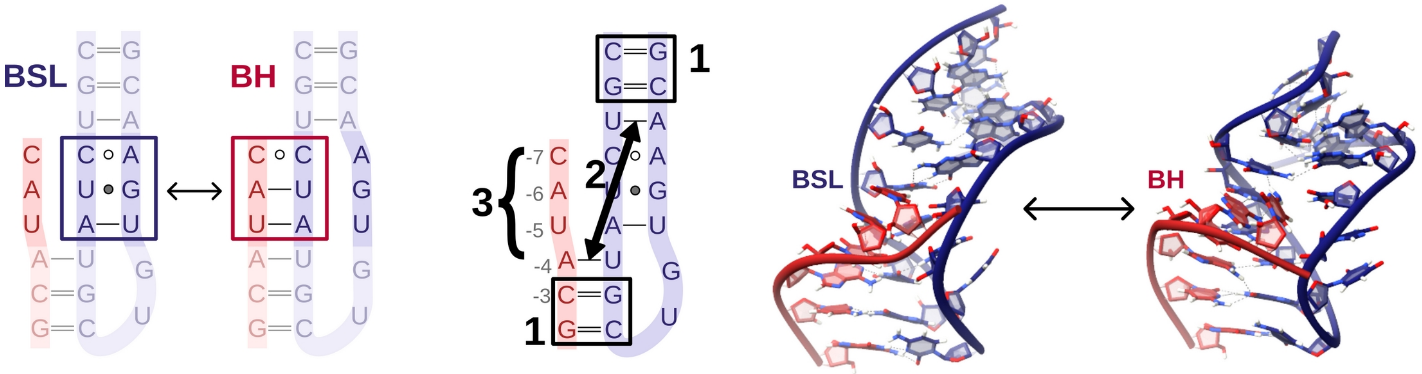

**Figure S1:** RNA model system used in the metadynamics simulations. The left panel shows a cartoon of the model with the intron sequence depicted in red and the BSL in blue. The box denotes the base pairs biased in metadynamics simulations. The middle panel presents a scheme of some of the applied wall restrains; numbering corresponds to the list reported in Supplementary results 1. The right panel shows the structures of the target BSL and branch helix (BH) states.

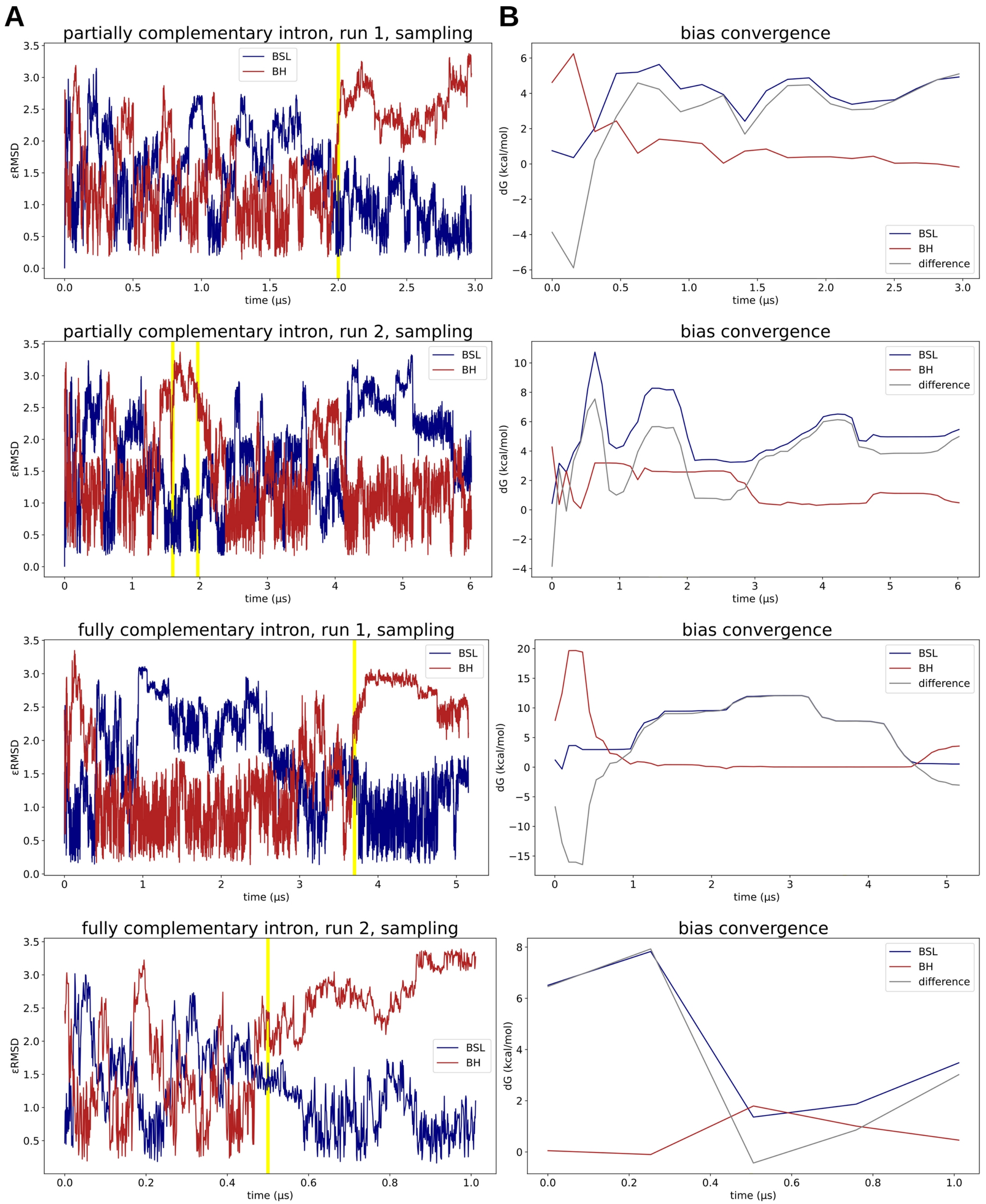

**Figure S2: Development of metadynamics simulations. (A)** εRMSD as function of simulation time for the different systems and replicas. The blue and the read lines report the εRMSD evolution with respect to the BSL and the branch helix target structures. The yellow vertical lines mark the time after which (or the interval in which) the simulations were truncated for the analyses. **(B)** Convergence of the non-averaged bias versus simulation time for the different systems and replicas. Note the different time scales in each graph.

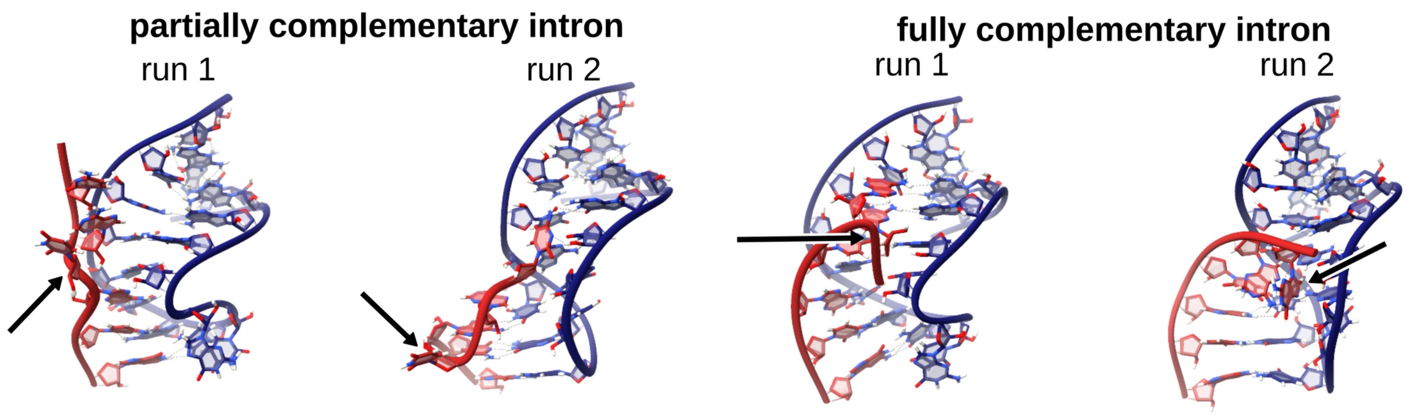

**Figure S3: Off-target states visited in metadynamics simulations of the intron (red ribbons) featuring partial (left) and full (right) base pair complementarity to the BSL (blue ribbons).** The arrows point to problematic interactions. Namely, in the first and second figures it is U-5 bulged out, in the third figure a sharp backbone turn caused G-7 stacking between A-6 and U-5, and in the last figure G-7 got stucked away from the BSL minor groove.

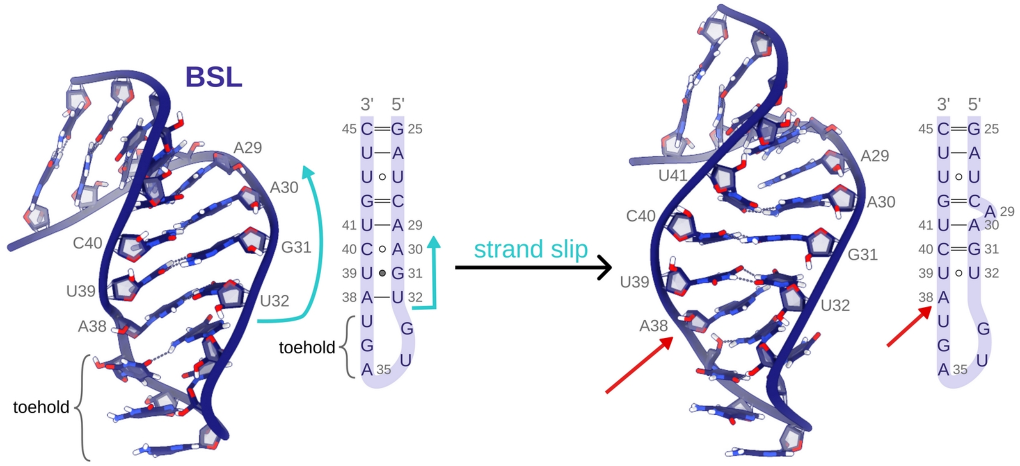

**Figure S4: Selected snapshots of MD simulation trajectories showing BSL (in blue cartoon or in sketch) strand slip leading to A38 remodeling suitable for intron binding**. U41 can bind to either A29 or A30. Introns are not shown for clarity. The two structures are aligned with respect to their toehold bases.

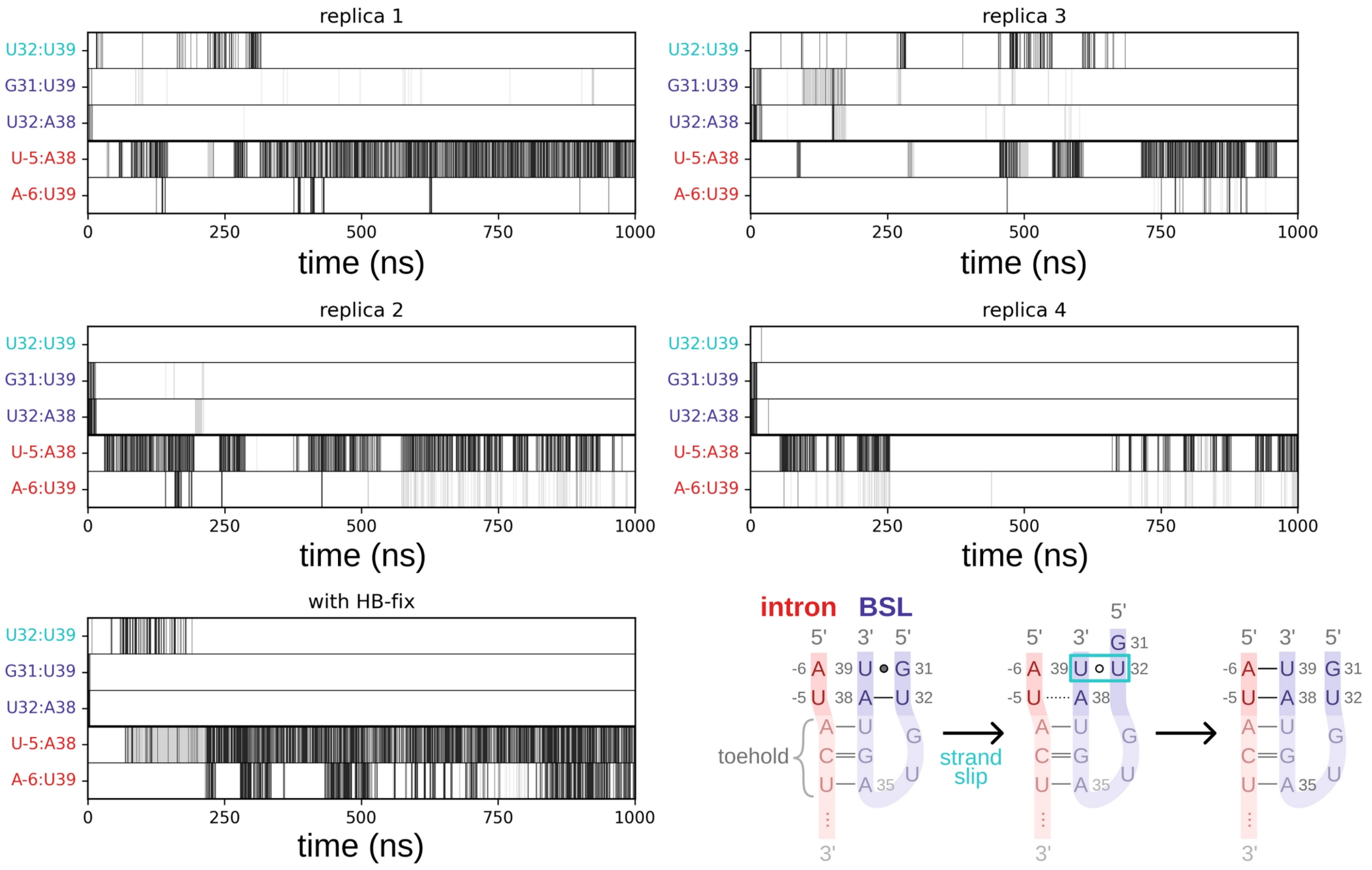

**Figure S5: MD simulations of the small intron-BSL construct.** Formation of the base pairs versus simulation time. Black, gray and white colors indicate the full formation of the respective pair (heavy-atom distance < 3.2 Å and angle > 140° for all expected H-bonds), weak pairing (distance < 3.5 Å and angle > 120° for at least one of the expected H-bonds), and the absence of the respective pairing. The hydrogen-bonds of BSL are labeled blue, while the hydrogen-bonds of the branch helix are red; the slip-stranded U32:U39 pair is cyan. Scheme of the simulated system is shown in the bottom right corner. The bases which are not analyzed in the graphs are shaded.

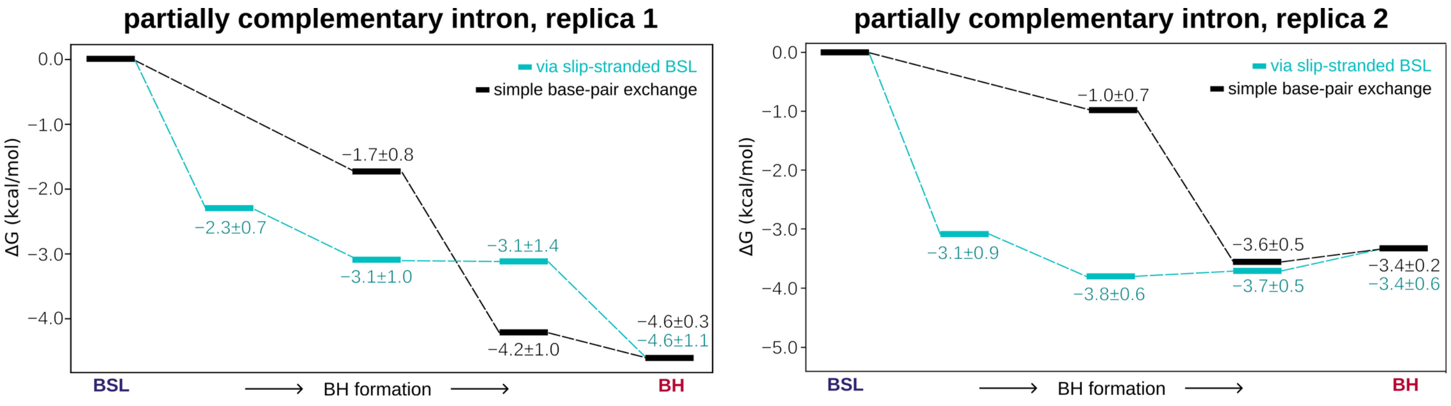

**Figure S6: Scheme of the relative free energy (kcal/mol) of the two competitive pathways (slip-stranded and base-pair exchange pathways) leading to the formation of the branch helix (BH).** The errors for an energy difference between the consecutive steps are reported. The slip-stranded state is the driving force for the slip stranded pathway, while the most significant stabilization base-pair exchange pathways on the black pathway is due to the formation of two new BH base pairs.

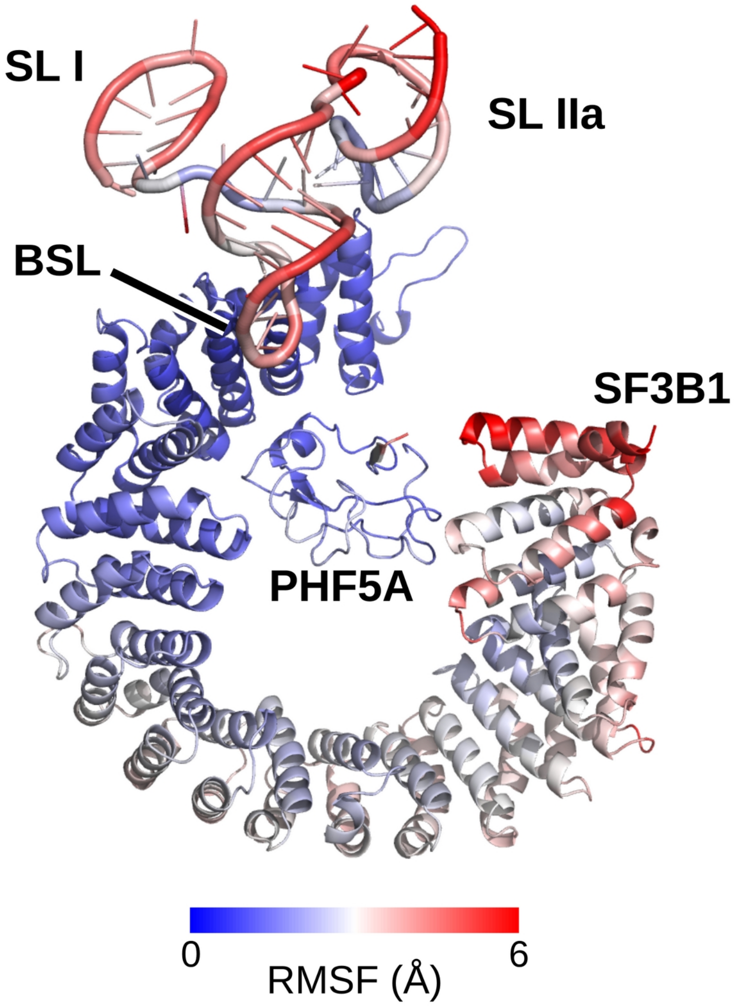

**Figure S7: Root-mean-square fluctuations (RMSF) calculated on the MD simulation trajectories of the U2/SF3b particle with TAT-SF1.** Trajectories of two replicas were collectively analyzed. Only the RNA, SF3B1, and PHF5A proteins are shown for clarity. The loop of SL IIa is anchored on the SF3B1 protein.

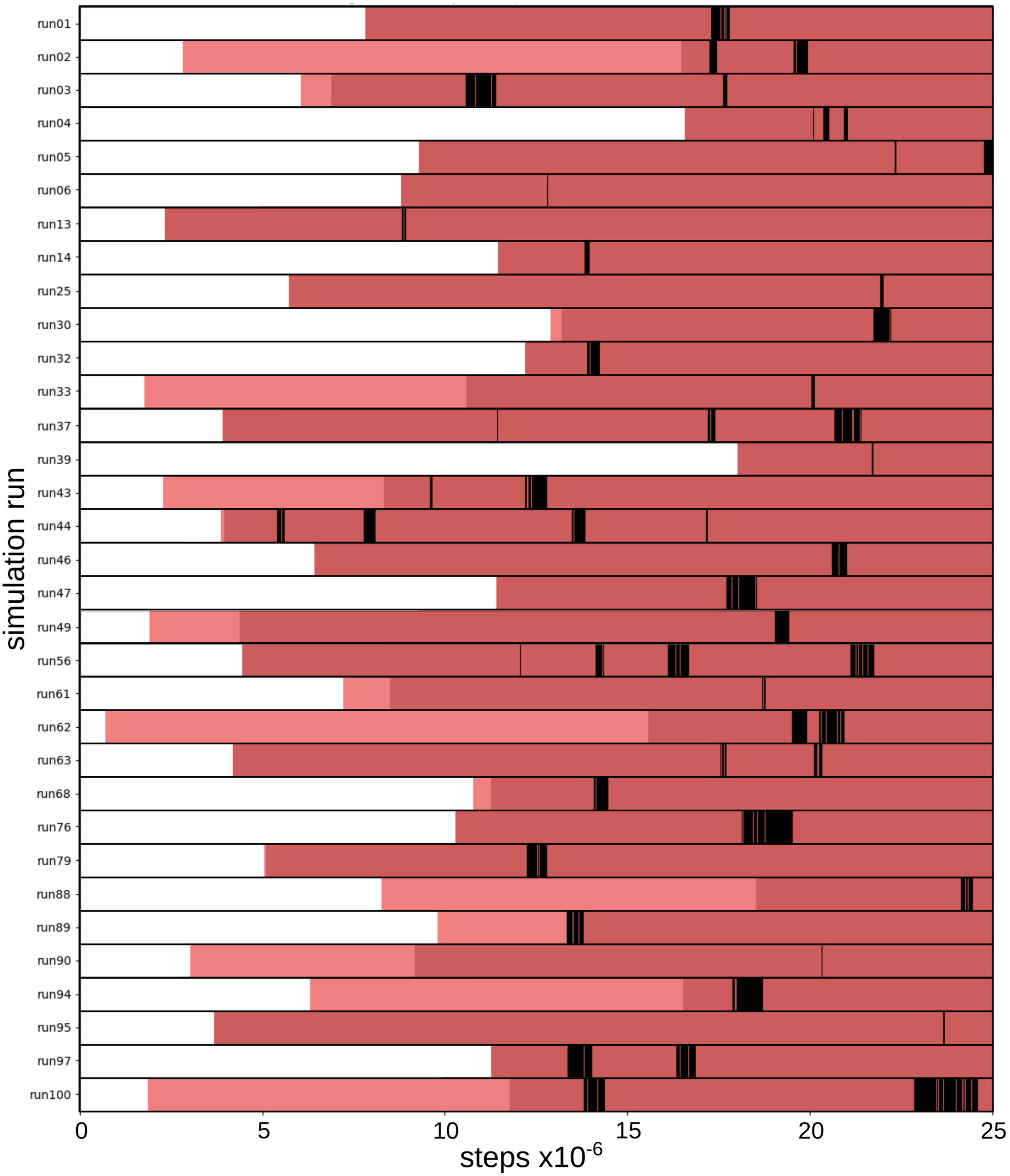

**Figure S8: Branch point adenosine (BPA) bulging out events vs simulation steps in coarse grained simulations.** Only the trajectories that sampled BPA bulge out events are shown. In the histogram, the light red marks the formation of the second last base pair of the branch helix end flanking the BPA (A3:U32), dark red marks the formation of the last pair (C4:G31), and the black areas mark the BPA bulge out events.

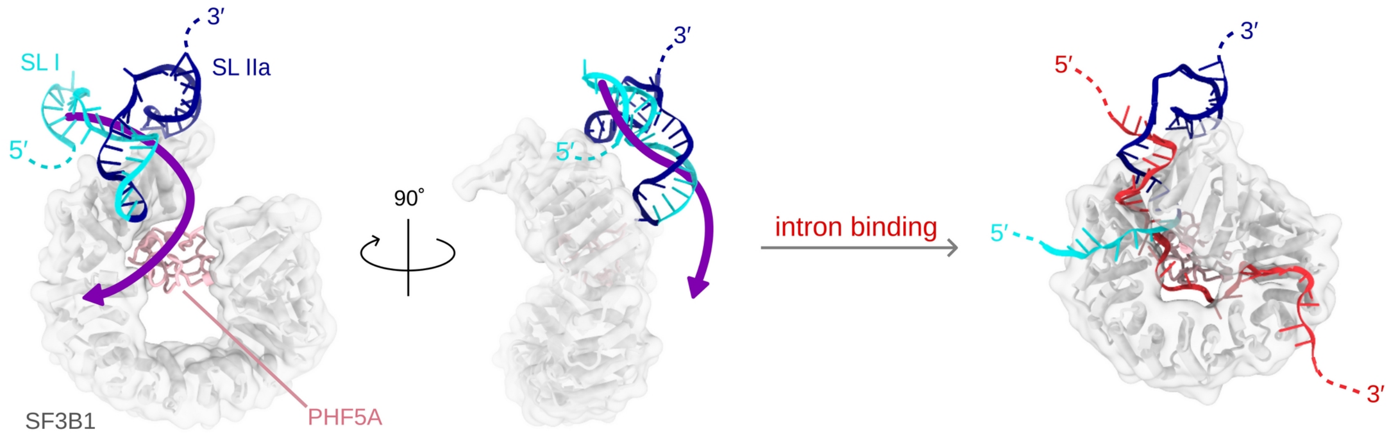

**Figure S9:** Relocation of U2 snRNA 5’-end (in light blue) necessary to create space for intron binding and formation of the branch point helix. The putative relocation direction is indicated by a violet arrow.
